# Extensive Evaluation of Weighted Ensemble Strategies for Calculating Rate Constants and Binding Affinities of Molecular Association/Dissociation Processes

**DOI:** 10.1101/671172

**Authors:** A. J. Pratt, Ernesto Suárez, Daniel M. Zuckerman, Lillian T. Chong

## Abstract

The weighted ensemble (WE) path sampling strategy is highly efficient in generating pathways and rate constants for rare events using atomistic molecular dynamics simulations. Here we extensively evaluated the impact of several advances to the WE strategy on the efficiency of computing association and dissociation rate constants (k_on_, k_off_) as well as binding affinities (K_D_) for a set of benchmark systems, listed in order of increasing timescales of molecular association/dissociation processes: methane/methane, Na^+^/Cl^-^, and K^+^/18-crown-6 ether. In particular, we assessed the advantages of carrying out (i) a large set of “light-weight” WE simulations that each consist of a small number of trajectories vs. a single “heavy-weight” WE simulation that consists of a relatively large number of trajectories, (ii) equilibrium vs. steady-state WE simulations, (iii) history augmented Markov State Model (haMSM) post-simulation analysis of equilibrium sets of trajectories, and (iv) tracking of trajectory history (the state last visited) during the dynamics propagation of equilibrium WE simulations. Provided that state definitions are known in advance, our results reveal that heavy-weight, steady-state WE simulations are the most efficient protocol for calculating k_on_, k_off_, and K_D_ values. If states are not strictly defined in advance, heavy-weight, equilibrium WE simulations are the most efficient protocol. This efficiency can be further improved with the inclusion of trajectory history during dynamics propagation. In addition, applying the haMSM post-simulation analysis enhances the efficiency of both steady-state and equilibrium WE simulations. Recommendations of appropriate WE protocols are made according to the goals of the simulations (*e.g*. to efficiently calculate rate constants and/or generate a diverse set of pathways).

## 1. INTRODUCTION

The WE strategy^1-2^ has been demonstrated to be highly efficient relative to standard “brute-force” simulations in generating pathways and rate constants for host-guest association,^3^ protein-small molecule dissociation,^4-5^ and protein-peptide association^6^ using atomistic molecular dynamics (MD) simulations. While these applications demonstrate the power of the WE strategy, the development of the most efficient WE protocol has typically been done by trial-and-error given that no systematic comparison of efficiencies in calculating observables of interest has been completed – until now.

To aid users in identifying the most efficient WE protocols for their simulation goals, we evaluated several advances to the WE strategy in terms of their impacts on the efficiency of the WE strategy in calculating k_on_, k_off_, and K_D_ values for the following benchmark systems, listed in order of increasing structural complexity and free energy barrier for the molecular association/dissociation process: methane/methane (CH_4_/CH_4_), Na^+^/Cl^-^, and K^+^/18-crown-6 ether (K^+^/CE) (Fig. 1). In particular, we assessed the advantages of carrying out the following in conjunction with explicit-solvent MD simulations: (i) equilibrium vs. steady-state WE simulations, (ii) a large set of “light-weight” WE simulations that each consist of a small number of trajectories vs. a single “heavy-weight” WE simulation that consists of a relatively large number of trajectories, (iii) haMSM post-simulation analysis, and (iv) tracking of the “history”, or state last visited by each trajectory, during dynamics propagation in equilibrium WE simulations. The ultimate goal of our evaluation was to determine the most efficient WE simulation protocol for calculating the k_on_, k_off_, and K_D_ values for receptor-ligand association/dissociation processes that are much slower than the microsecond timescale and therefore inaccessible to brute force simulations. Such observables have been of great interest to a variety of research areas in chemistry and biology, including host-guest interactions, protein engineering, and drug discovery. Furthermore, the generation of complete pathways for such processes at the atomic level can be highly valuable for aiding efforts to improve the kinetics of receptor-ligand interactions by providing structures of transient states.

**FIG. 1.**
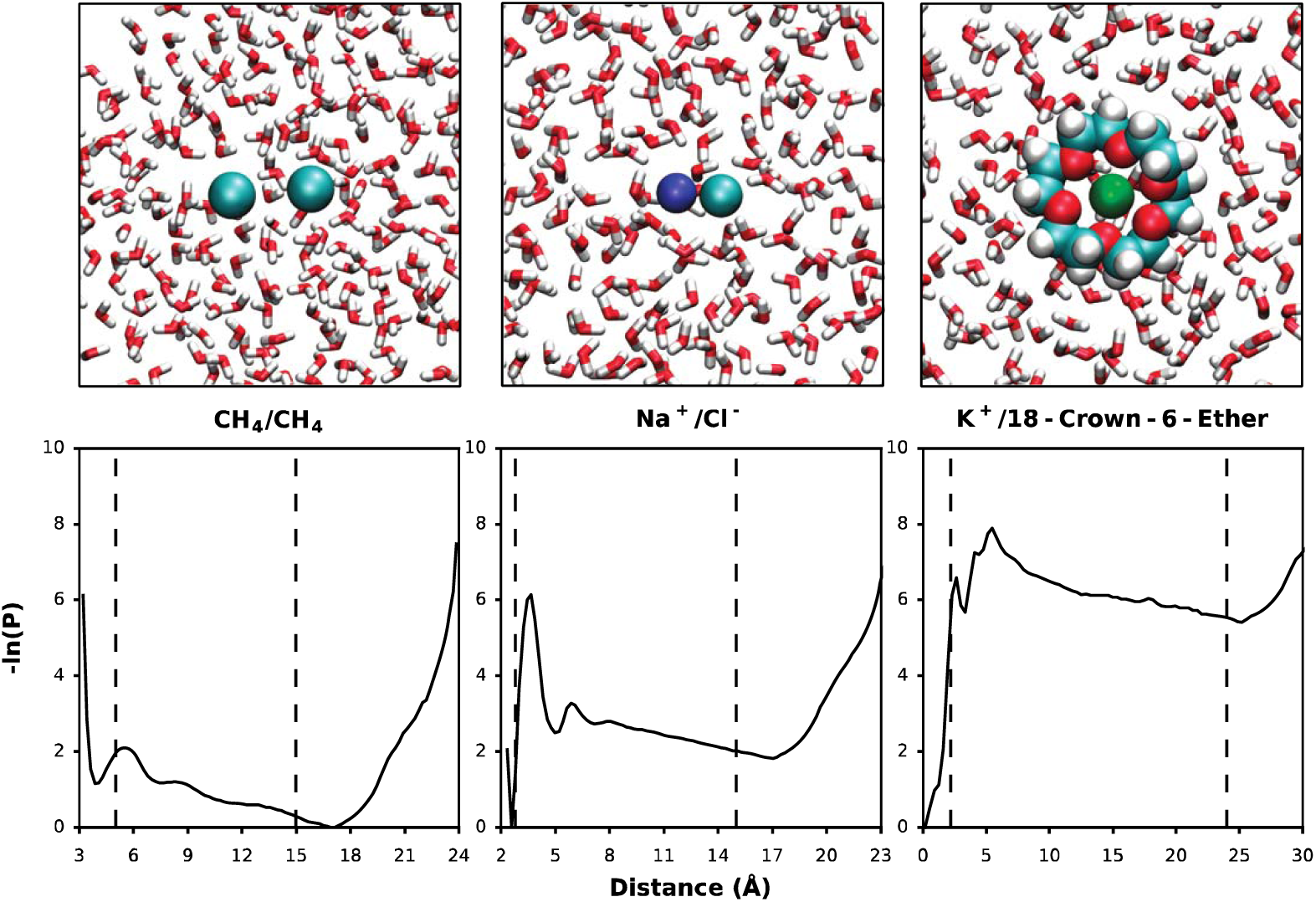
Benchmark systems of study. Systems are immersed in explicit water and displayed in order of increasing structural complexity and barrier to the molecular association/dissociation process, as estimated by probability distributions as a function of the relevant progress coordinate (see Methods) from brute force simulations. Dashed lines indicate boundaries used for definitions of unbound and bound states.

## 2. THEORY

The essence of the weighted ensemble (WE) path sampling strategy is to carry out many trajectories in parallel, with each trajectory assigned a weight to properly represent the ensemble. To control the trajectory distribution, configurational space is divided into bins along a progress coordinate for tracking trajectory progress towards the target state. Trajectories are examined at fixed time intervals τ and evaluated for replication or pruning to ensure even coverage of configurational space with a target number of trajectories/bin. During this resampling procedure, trajectory weights are adjusted according to rigorous statistical rules such that no bias is introduced into the dynamics. The result of the procedure is a “statistical ratcheting” effect that enhances the efficiency of generating continuous pathways and rate constants for the rare event of interest. Features of the original WE strategy along with recent advances that are evaluated in this study are presented below.

### 2.1 Equilibrium vs. steady-state WE simulations

WE simulations can be carried out under steady-state or equilibrium conditions. In the original WE strategy by Huber and Kim^1^, steady-state conditions are maintained by “recycling” trajectories that have reached the target state, *i.e*. starting new trajectories, with the same weights from the initial state. Recent advances to the strategy^2^ have enabled WE simulations to be run under equilibrium conditions, which does not require recycling trajectories once they reach the target state. Thus, a significant advantage of such equilibrium WE simulations is that *states do not need to be strictly defined in advance*, but can instead be approximately defined during dynamics propagation and refined after the completion of the simulation. In addition to providing equilibrium observables (*e.g*. state populations), equilibrium WE simulations can be divided into two steady state ensembles of trajectories based on the “history”‘ label, or state last visited, to enable the direct calculation of rate constants corresponding to each steady state^3^.

### 2.2 History augmented Markov State Model (haMSM) post-simulation analysis

To enable the calculation of long-timescale observables from shorter simulations under equilibrium conditions, the history augmented Markov State Model (haMSM; also referred to previously as non-Markovian analysis) post-simulation analysis can be applied after generating a set of equilibrium trajectories^7^ either via equilibrium WE simulation or the combination of steady-state WE simulations in opposite directions (association and dissociation directions). In this procedure, the resulting equilibrium set of trajectories is first decomposed into subsets of trajectories that correspond to each of the two steady states. Next, a rate matrix (or transition probability matrix) is constructed from history-labeled transition probabilities 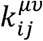 between bins (*i* and *j*) for trajectories initially in the *µ* subensemble which transition to the *ν* subensemble, where *µ* and *ν* are either of the two steady states *α* and *β*. This labeled rate matrix is then used to solve the set of standard steady-state equations to yield the populations of each bin *i* corresponding to the two steady states *α* and *β* (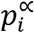and 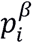) and ultimately, the equilibrium population of bin 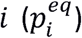, which is the sum of the two steady-state populations in that bin 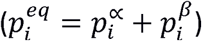. The haMSM analysis procedure enables the calculation of equilibrium as well as non-equilibrium observables from an equilibrium set of trajectories without a Markovian assumption. Furthermore, the haMSM procedure can be carried out using any arbitrary set of bins and history labels and is therefore not restricted to those used during dynamics propagation. Thus, for each benchmark system in the present study, we tested the use of the minimal set of bins (*i.e*. three bins for delineating the initial state, transition region, and target state; see Fig. 1) as well as larger sets of bins.

### 2.3 Tracking the state last visited (“history”) during dynamics propagation

To ensure that pathways are generated in both directions (molecular association and disassociation) in an equilibrium WE simulation, the trajectory history (state last visited) can be included as a dimension of the progress coordinate. A potential issue with equilibrium WE simulations is that a bin may contain trajectories corresponding to each of the two steady state ensembles, and trajectories that correspond to one of the steady states may be pruned. Tracking the history of each trajectory as a dimension of the progress coordinate separates the trajectories into the corresponding steady-state ensembles during the dynamics propagation and avoids over-pruning trajectories in one steady state.

### 2.4 Light-weight vs. heavy-weight WE simulations

Two alternate strategies for carrying out WE simulations are (i) running a single “heavy-weight” simulation that consists of a large target number of trajectories/bin (50 in the present study), and (ii) running a large set (50 in the present study) of “light-weight” simulations that each consist of a relatively small number of target trajectories/bin (2 or 4 in the present study). While the statistical ratcheting effect would be expected to be greater for heavy-weight vs. light-weight simulations due to the larger number of target trajectories/bin, many of the successful pathways that are generated by heavy-weight simulations will be correlated in history, *i.e*. sharing common trajectory segments, which can degrade the statistical quality of the data. An advantage to carrying out a large set of light-weight simulations over a single heavy-weight simulation is that the set of light-weight simulations may yield a greater diversity of pathways that do not share trajectory history and are independent, which enables more straightforward calculation of error in the observables of interest.

## 3. METHODS

To extensively evaluate the advances to the WE strategy mentioned above, we carried out 10 simulation protocols for each of the three benchmark systems (Table 1), yielding a total of 304 WE simulations of molecular dissociation and/or association processes for each system with an aggregate simulation time of 5.67, 47.8, and 29.7 µs for the CH_4_/CH_4_, Na^+^/Cl^-^, and K^+^/CE systems, respectively. In all of our comparisons of the various WE protocols, the number of target trajectories/bin for each steady-state set of trajectories was chosen to be approximately equal to maintain a similar level of statistical ratcheting. For example, single equilibrium, heavy-weight simulations with 50 target trajectories/bin were compared with pairs of steady-state, heavy-weight simulations with 50 target trajectories/bin. For each benchmark system, an unprecedented amount of sampling of molecular association/dissociation events was achieved using 912 WE simulations and an aggregate simulation time of 83.17 μs within one month using 288 Intel 2.6 GHz CPU cores at a time.

**Table 1:** WE simulation protocols tested in this study. For each of the three benchmark systems, the 10 simulation protocols listed below were tested, comparing (i) simulations under steady state vs. equilibrium conditions, and (ii) the use of a single heavy-weight simulation vs. a set of 50 light-weight simulations. Steady-state WE simulations were started from both bound and unbound states while equilibrium WE simulations were started only from bound states. In addition, we assessed the effects of tracking the trajectory history during dynamics propagation on the efficiency of equilibrium WE simulations in calculating observables of interest.

| steady-state/<br>equilibrium | heavy-weight/<br>light-weight | # WE<br>simulations | # target<br>trajectories/bin | initial<br>state | history<br>tracking? |
| --- | --- | --- | --- | --- | --- |
| steady state | heavy-weight | 3 | 50 | bound | -- |
|  |  | 3 | 50 | unbound | -- |
|  | light-weight | 50 | 2 | bound | -- |
|  |  | 50 | 2 | unbound | -- |
|  |  | 50 | 4 | bound | -- |
|  |  | 50 | 4 | unbound | -- |
| equilibrium | heavy-weight | 3 | 50 | bound | Y |
|  |  | 1 | 100 | bound | N |
|  | light-weight | 50 | 2 | bound | Y |
|  |  | 50 | 4 | bound | Y |

### 3.2 WE simulations

All WE simulations were carried out using the open-source, highly scalable WESTPA software package^8^. To ensure that the starting structures for our WE simulations were representative of the corresponding state (bound or unbound states), we sampled the initial state under equilibrium conditions (Table S1 in supporting information). The resulting ensembles served as a pool of starting structures that could be selected according to their statistical weights for running separate WE simulations of the molecular dissociation or association processes. All equilibrium WE simulations were started from a pre-equilibrated ensemble of bound-state structures, whereas steady-state WE simulations were started from a pre-equilibrated ensemble of either the unbound- or bound-state structures to generate molecular association or dissociation pathways, respectively.

For each benchmark system, the same progress coordinates as those used in a previous WE study ^3^ were used for the WE simulations: i) for CH_4_/CH_4_, the distance between the carbon atoms of the two CH_4_ molecules, (ii) for Na^+^/Cl^-^, the distance between the two ions, and (iii) for K^+^/CE, the distance between the K^+^ ion and the center-of-mass of the crown ether oxygens. While the focus of this study was not to systematically examine bin choices, a few refinements were made to the previously defined bins (Table S3 in Supporting Information). First, bin positions were adjusted along with the inclusion of additional bins to further enhance the sampling of the dissociation processes. Second, state definitions, particularly those of the unbound state, were refined for both the CH_4_ /CH_4_ and K^+^/CE systems to be more stringent in order to emcompass only the region of the basin thereby yielding more consistent rate constants among a set of independent WE simulations. For the Na^+^/Cl^-^ system, the same state definitions as those in the previous WE study^3^ were used. Finally, the history of each trajectory (state last visited) was tracked during dynamics propagation for a subset of the equilibrium WE simulations by including history as a second dimension of the progress coordinate, using the same state definitions.

### 3.2 Dynamics propagation

As done previously,^3^ dynamics were propagated for the CH_4_ /CH_4_ and Na^+^/Cl^-^ systems using the GROMOS 45A3 force field^9^ and SPC/E water model,^10^ and for the K^+^/CE system, using the OPLS-AA/L force field^11^ and TIP3P water model.^12^ All of our simulations included a minimum 12-Å clearance between the solutes in their unbound states and the box walls.

### 3.3 Standard “brute-force” simulations

Extensive standard “brute force” simulations of the molecular association/dissociation processes of the CH_4_/CH_4_, Na^+^/Cl^-^, and K^+^/CE systems were generated on the µs-timescale by our previous study.^3^

### 3.4 Evaluation of simulation convergence

A WE simulation was considered converged if the instantaneous mean of the observable of interest at the final iteration of the simulation was (i) within the 95% confidence interval of the mean values at prior iterations, and (ii) within the 95% confidence interval of the mean values from other simulation protocols (Figs. S1 and S2). According to these criteria, all of our simulations were sufficiently long to achieve convergence.

### 3.5 Application of the haMSM post-simulation analysis

For each benchmark system, the haMSM post-simulation analysis^7^ was applied to a set of equilibrium trajectories using five sets of bins: the minimal set of 3 bins representing the unbound state, transition region, and bound state (Table S2 in Supporting Information), and four additional sets of 4, 6, 10, and 18 bins, as generated by successively halving the bins in the transition region. For each set of bins and each WE iteration, a history-labeled transition probability matrix was constructed from a running average of bin-to-bin fluxes and bin populations for the same iteration of each set of equilibrium trajectories (equilibrium WE simulations or combinations of steady-state WE simulations in opposite directions), and then solved to obtain steady-state fluxes *f*_*ij*_ from states *i* to *j* and steady-state populations of the unbound and bound states. As mentioned in section 2.2, equilibrium state populations were estimated as the sum of each steady-state population in the corresponding bins.

### 3.6 Calculation of rate constants

The same strategy was used to calculate rate constants for both WE and brute force simulations. The rate constant k_ij_ for transitions from bin *i* to *j* was calculated using the following ratio of averages:^7^

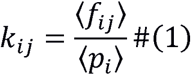

where ⟨*f*_*ij*_ ⟩ is the running average of the flux from bin *i* to *j* of the steady-state set of trajectories in the direction of interest (e.g., bound to unbound) and ⟨*p*_*i*_ ⟩ is the running average of the steady-state population of the initial state *i* across all iterations *t* with a fixed length *τ*. Note that for steady-state WE simulations and equilibrium WE simulations with very few events in the reverse direction, *p_i_* is unity.^6^ For equilibrium WE simulations with a significant number of events in the reverse direction, *k*_*ij*_ is calculated by normalizing the average *f*_*ij*_ by the average *p*_*i*_ thereby focusing the calculation only on the unidirectional flux that corresponds to the steady state of interest.

### 3.7 Calculation of binding affinities (K_D_ values)

Binding affinities (K_D_ values) were calculated using two different approaches. In the first approach, equation (2), we calculated K_D_ values from the kinetic observables, *i.e.* the k_off_ and k_on_:

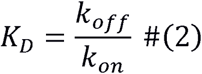

In the second approach, equation (3), we calculated K_D_ values from equilibrium observables, *i.e*. equilibrium state populations of the unbound state (*p*_*U*_ and bound state (*p*_*B*_):

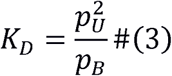

Equilibrium state populations were calculated as the steady-state population of that state by applying the haMSM post-simulation analysis to both brute force and WE simulations. Uncertainties were calculated using error propagation from the 95% confidence intervals of the rate constants or state populations.

### 3.8 Estimation of WE efficiency in computing rate constants and binding affinities

The efficiency *S*_*k*_ of a given WE protocol in computing the rate constant of interest (k_on_ or k_off_) was estimated using the following equation:^1^

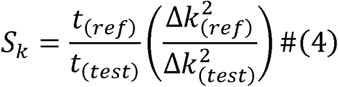

where *t*_(*ref*/*test*)_ is the aggregate simulation time for the reference or test simulation, respectively, and Δ*k*_(*ref*/*test*)_ is the relative error in the rate constants (uncertainty of the rate constant relative to the rate constant where the uncertainty represents the 95% confidence interval) for the corresponding simulations. Thus, the efficiency of a test simulation in calculating the rate constant is determined by taking the ratio of aggregate times for the test and reference simulations that would be required to estimate the rate constant with the same relative error, assuming that the square of the width of the 95% confidence interval on the rate constant is inversely proportional to the simulation time.^1^

Similarly, the efficiency *S*_*k*_ of computing the K_D_ was estimated using the following equation (equation (6)):

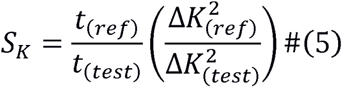

where Δ*K*_*(ref/test)*_ is the relative error in K_D_. In cases where the K_D_ was calculated from the ratio of rate constants, the corresponding *S*_*k*_ is also a metric of how efficient a simulation is in computing the rate constants in both directions.

For both WE and brute force simulations, the relative errors of the observables of interest were calculated using the same error estimation strategy. In particular, a Monte Carlo bootstrap procedure^13^ was applied to estimate the running average and uncertainties of rate constants and state populations from each simulation set. These uncertainties were then propagated to estimate the relative errors in the in the K_D_ values using either equations (2) or (3). Complete details of the bootstrap procedure, including differences in its application to direct calculation vs. haMSM post-analysis of the rate constants and state populations are summarized in Table 2. For each WE protocol, estimates of the efficiency *S*_*k*_ were based on the final iterations. As mentioned in section 3.4, all WE simulations were sufficiently long to satisfy the convergence criteria.

### 3.9 Estimation of sampling variance for simulation protocols

To assess how a simulation protocol affects the efficiency of calculating rate constants, a novel (to our knowledge) network analysis was conducted. In particular, Sampling Error Networks (SEN) were constructed to visualize the precision in sampling bin to bin transitions. Transition matrices, generated from each simulation replicate, were used to estimate the variance in sampling a given bin to bin transition as the error in the simulation observables is ultimately related to the variance in the transition matrices. Each node in the SEN represents a bin used during the simulation, and the log of the inverse of the percent error was used as the weight for the edges connecting nodes. A force directed layout algorithm in Gephi, ForceAtlas2, was used to generate the layout. A bin to bin transition was sampled if two bins are connected, and their distance is positively correlated to the sampling variance.

## 4 RESULTS AND DISCUSSION

We extensively evaluated the impact of several advances in the WE strategy on the efficiency of calculating the k_on_, k_off_, and K_D_ for the molecular association and dissociation processes of the following benchmark systems, listed in order of increasing timescales of the association/dissociation processes: methane/methane (CH_4_/CH_4_), Na^+^/Cl^-^, and K^+^/18-crown-6 ether (K^+^/CE). In particular, we determined the impact of the following advances on the efficiencies of the WE strategy in calculating the observables of interest: (i) a single heavy-weight simulation vs. a large set of light-weight simulations, (ii) steady-state vs. equilibrium WE simulations, (iii) tracking of trajectory history of during the dynamics propagation of equilibrium WE simulations, and (iv) history augmented Markov State Model (haMSM) post-simulation analysis^7^ of an equilibrium set of trajectories, generated by a single equilibrium WE simulations or pairs of steady state WE simulations. In the present study, equilibrium observables (*i.e*. the K_D_ from the ratio of state populations) were only calculated using the application of the haMSM analysis to an equilibrium set of trajectories (generated using a single equilibrium WE simulation or pair of steady state WE simulations) as the relevant equilibrium state populations did not otherwise converge.

### 4.1 Steady-State WE: Heavy-weight vs. Light-weight

We begin by comparing the efficiencies of carrying out WE simulations under steady-state conditions using a single heavy-weight simulation with a target number of 50 trajectories/bin vs. a set of 50 light-weight simulations using a target number of either 2 or 4 trajectories/bin. As mentioned above, equilibrium observables (i.e. the K_D_ from equilibrium state populations) can be calculated by combining trajectories from steady-state WE simulations in opposite directions to generate an equilibrium set of trajectories.

As shown in Fig. 2, the efficiency in calculating the k_on_, k_off_, and K_D_ (both from rate constants and equilibrium state populations) for all of the benchmark systems increased with the number of target trajectories/bin, with single heavy-weight simulations being the most efficient. This increase in efficiency is likely due to the greater extent of statistical ratcheting that would be expected to result from a greater number of chances to advance toward the target state (unbound or bound state). It is worth noting that the minimal number of target trajectories/bin (*i.e.* 2 trajectories/bin) is less efficient than brute force simulations in computing the k_on_ and K_D_ from the ratio of rate constants for the CH_4_/CH_4_ and Na^+^/Cl^-^ systems, and that a minimum of 4 target trajectories/bin is required to achieve greater efficiency than brute force simulations in computing these observables. Consistent with previous WE studies,^3, 14-17^ the efficiency of each type of WE simulation in calculating all of the observables increases with the height of the relevant free energy barrier, *e.g.* relative to brute force simulations, the efficiencies of single heavy-weight simulations in calculating the k_off_ are 2-to 5-fold for the CH_4_/CH_4_ system, 5-to 13-fold for the Na^+^/Cl-system, and 36- to 170-fold for the K^+^/CE system.

**FIG. 2.**
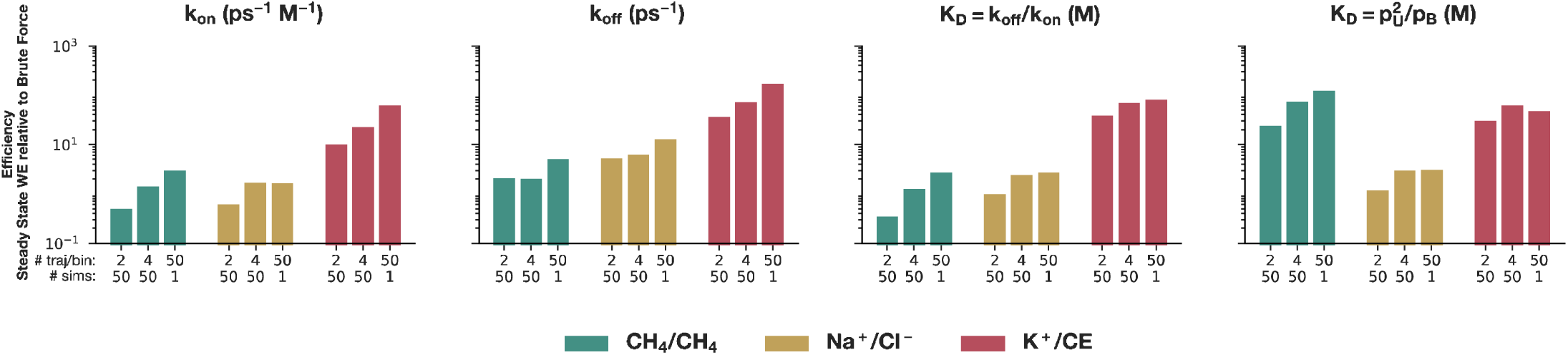
Efficiency of steady-state WE simulations relative to brute force simulations in calculating the k_on_, k_off_, and K_D_ for each benchmark system. The k_on_ and k_off_ values were calculated using steady-state WE simulations started from the unbound and bound states, respectively. The K_D_ values were calculated using pairs of steady-state WE simulations in opposite directions; the K_D_ was evaluated using both the rate constants (k_off_/k_on_) and equilibrium populations for the unbound state (p_U_) and bound state (p_B_); the haMSM post-simulation analysis was applied in the latter case. Data is shown for (i) sets of 50 light-weight simulations with 2 and 4 target trajectories/bin, and (ii) a single heavy-weight simulation with 50 target trajectories/bin. The horizontal gray line indicates equal efficiency relative to brute force simulations.

### 4.2 Equilibrium WE: Heavy-weight vs. Light-weight

Next, we compared the efficiencies of carrying out WE simulations under equilibrium conditions using a single heavy-weight simulation vs. 50 light-weight simulations with either 2 or 4 target trajectories/bin). As mentioned above, equilibrium WE simulations can be decomposed into two subsets of steady state trajectories for the molecular association and dissociation directions, and can therefore yield k_on_ and k_off_ as well as K_D_ values. As shown in Fig. 3, the efficiencies of such simulations in calculating the k_on_, k_off_, and K_D_ generally increased with the number of target trajectories/bin, as found above for the steady state WE simulations. An exception to this trend was the calculation of the k_on_ for the Na^+^/Cl^-^ system for which a single heavy-weight simulation was less efficient than 50 light-weight simulations with 4 trajectories/bin, but more efficient than 50 light-weight simulations with 2 trajectories/bin. Nonetheless, our results indicate that in general, heavy-weight simulations under both steady-state and equilibrium conditions are the most efficient simulation protocol for calculating rate constants and binding affinities.

**FIG. 3.**
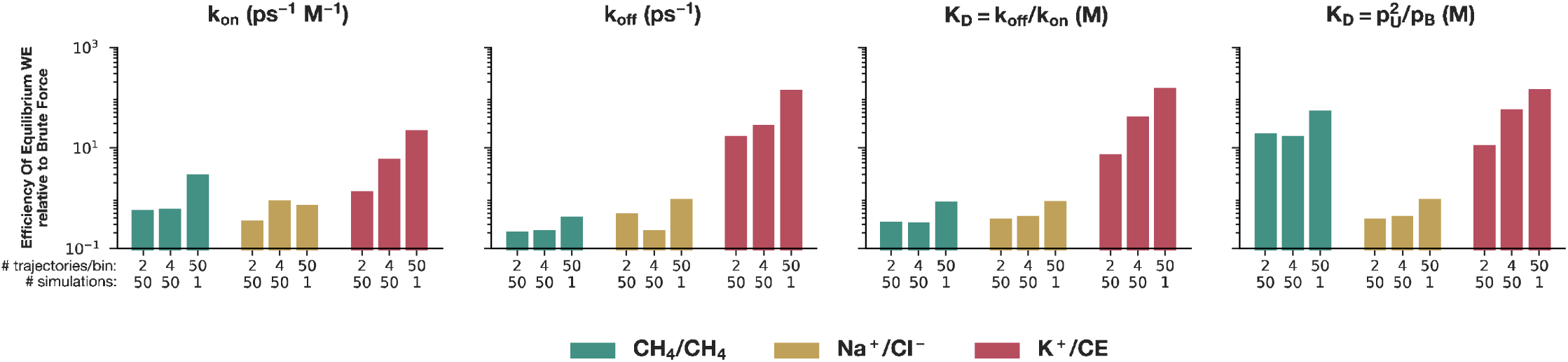
Efficiency of equilibrium WE simulations relative to brute force simulations in calculating the k_on_, k_off_, and K_D_ for each benchmark system. The K_D_ values were calculated using both the rate constants (k_off_/k_on_) and equilibrium populations for the unbound state (p_U_) and bound state (p_B_). The latter involved application of the haMSM post-simulation analysis. Data is shown for (i) sets of 50 light-weight simulations with 2 and 4 target trajectories/bin, and (ii) a single heavy-weight simulation with 50 target trajectories/bin. The horizontal gray line indicates equal efficiency relative to brute force simulations.

### 4.3 Equilibrium vs. Steady-State WE

We then compared the efficiency of carrying out steady-state WE simulations vs. equilibrium WE simulations in calculating the k_on_, k_off_, and K_D_ for each of the benchmark systems. To calculate the K_D_ values, pairs of steady-state WE simulations in opposite directions were compared with single equilibrium WE simulations. In calculating the k_on_ and k_off_, it is not surprising that equilibrium WE simulations are less efficient than steady state WE simulations with the appropriate target state (bound or unbound states, respectively) since the former involve the use of additional computational effort to generate pathways in the opposite direction from the target state (Fig. 4). Furthermore, despite the advantage of being able to simultaneously sample trajectories in opposite directions, a single equilibrium WE simulation is generally less efficient in calculating K_D_ values than a pair of steady-state WE simulations in opposite directions. The exception is the most complicated system, K+/CE, for which single equilibrium WE simulations are more efficient than pairs of steady-state WE simulations in calculating the K_D_ when a heavy-weight protocol is used in both cases (50 target trajectories/bin).

**FIG. 4.**
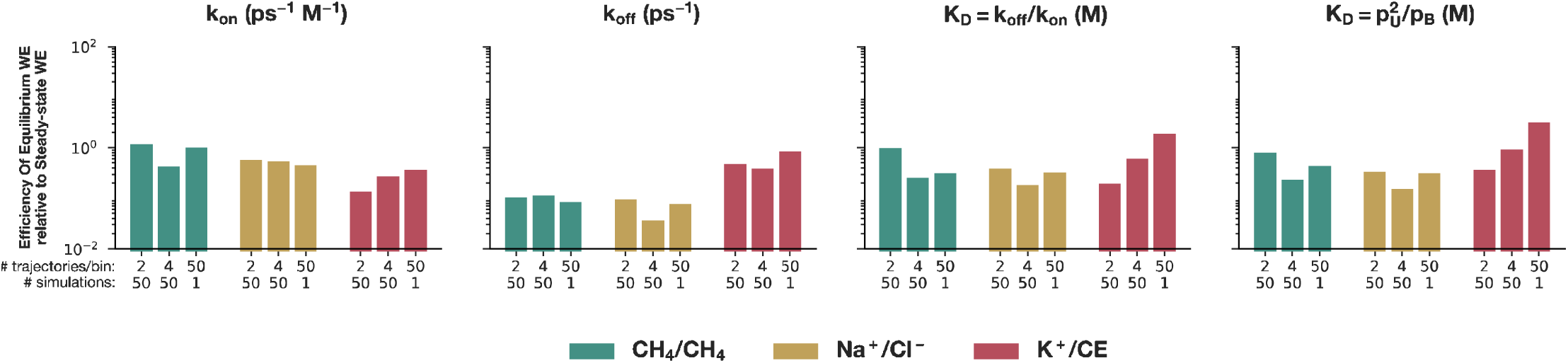
Efficiency of equilibrium WE simulations relative to the relevant steady state WE simulations in calculating the k_on_, k_off_, and K_D_ for each benchmark system. Relevant steady state WE simulations have the same number of WE simulations and target number of trajectories/bin (*e.g.* the efficiency of heavy-weight equilibrium simulations in calculating the K_D_ is determined relative to pairs of heavy-weight steady-state simulations in opposite directions). The horizontal gray line indicates equal efficiency relative to the relevant steady state WE simulations.

To determine the reason for the lower than expected sampling efficiencies for CH_4_/CH_4_ and Na^+^/Cl^-^ in the heavyweight steady state case, the simulation data was used to construct a series of sampling error network plots (Fig. 5). A SEN with tightly packed clusters and nodes corresponds to a simulation protocol that has less variance in sampling transitions than a simulation protocol with a SEN occupying more space.

**FIG. 5.**
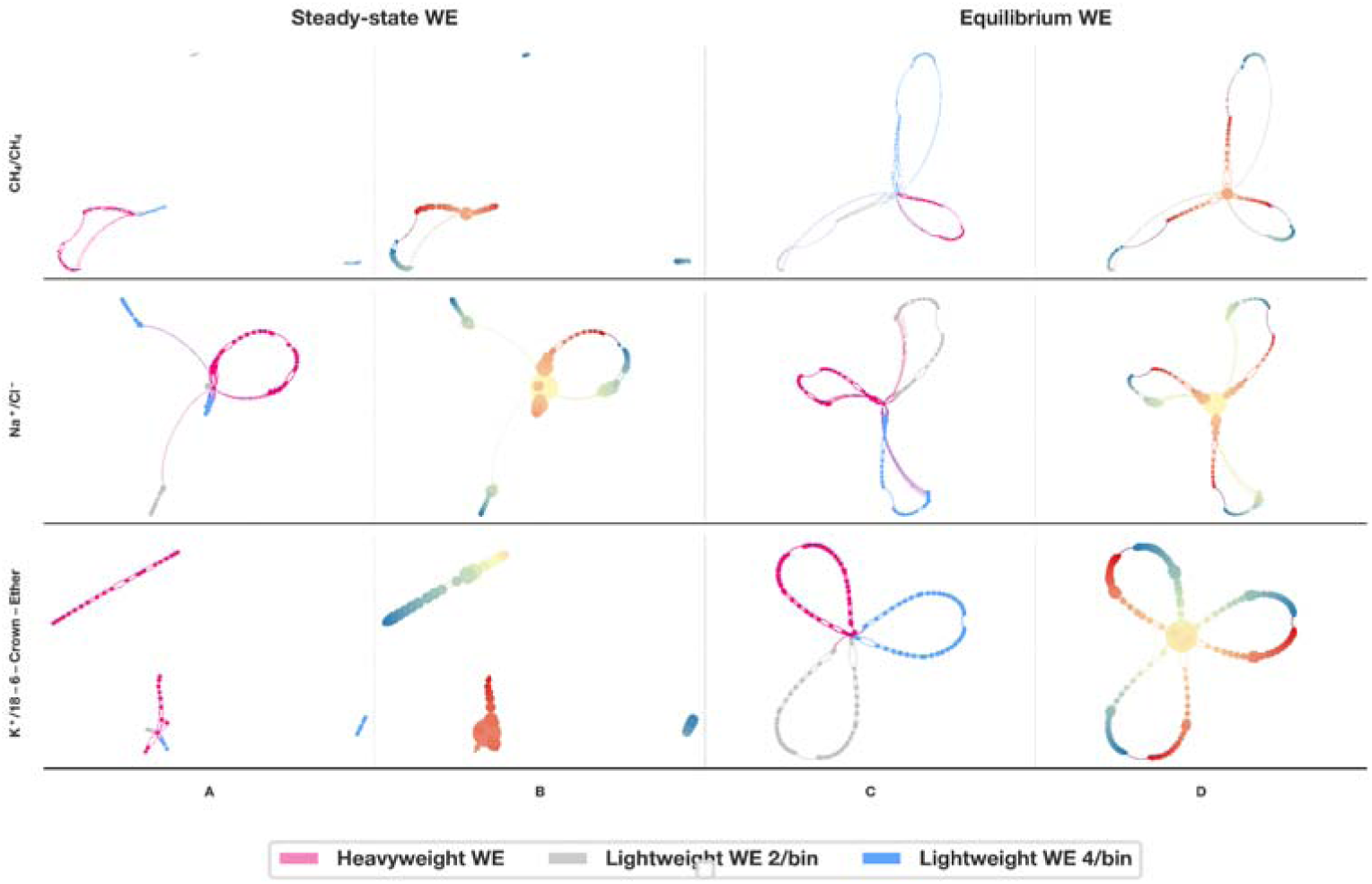
Sampling error networks (SEN) of simulation protocols. Each network contains 3 separate simulation protocols: heavyweight WE, lightweight WE with 2 trajectories/bin, and lightweight WE with 4 trajectories/bin (COLORS). Each node represents a bin used during dynamics, and the edges connecting the nodes have a strength equal to the inverse of the percent error in sampling the bin to bin transition corresponding to that node. Complete ‘lobes’ indicate that a complete binding/unbinding cycle was sampled with that simulation protocol; broken lobes are indicative of less successful sampling. Columns A & B are based on the same steady state simulation data, with different visualization schemes applied. In column A, the colors refer to the type of simulation, and the node sizes are equal. In column B, the color intensity is the progress coordinate distance (with red showing bins in the unbinding ensemble, and blue in the binding), and the node size is the sum of the weighted degree of the node. Columns C & D are the same as A & B, except they are for simulations run under equilibrium conditions. As the edge weights are the log of the inverse error, small changes in node size correspond to large differences in sampling variance.

The sampling error networks (SEN) suggest that the lower-than-expected efficiencies for calculating rate constants with the steady-state simulation protocol for Na^+^/Cl^-^ and CH_4_/CH_4_ may be due to simulation time spent working on reverse trajectories (i.e., binding trajectories in an unbinding simulation). The light-weight steady-state SENs for Na^+^/Cl^-^ and CH_4_/CH_4_ are unconnected across the binding/unbinding threshold, showing that trajectories which reach their target state were successfully recycled. The heavy-weight steady-state SENs, however, are connected, showing that the reverse trajectory was sampled within those simulations. Reverse trajectories add to the aggregate molecular time spent simulating and do not contribute to lowering error in estimating rate constants, reducing sampling efficiency. A lower tau value, or fewer walkers per bin, would likely improve the efficiency of these simulations.

Overall, the sampling error networks mirror the efficiency trends seen in Fig. 3 and 4, where equilibrium, heavy-weight WE networks have clusters which are more densely packed than those in either light-weight WE networks.

It may also be beneficial to reduce sampling in regions just beyond steep landscapes, as most of the simulation protocols show less variance in sampling transitions in and out of regions just before and after steep regions in the energy landscape. Columns B & D in Fig. 5, where the node radius is proportional to the weighted degree, show the largest nodes in regions just beyond steep climbs, suggesting that the variance in sampling transitions into and out of these regions is low. As these regions show lower variance in sampling transitions, some of that computation time may be better spent on the more difficult regions. In addition, K+/CE shows the largest nodes at a separation distance of 10 A, which is within the diffusive region. This suggests that sampling diffusive type mechanisms is straightforward for WE, and that the number of walkers in the diffusive region can be safely reduced.

### 4.4 Effects of haMSM post-simulation analysis

We then tested whether a haMSM post-simulation analysis of the same equilibrium sets of trajectories generated above would further increase the efficiency of calculating the k_on_, k_off_, and K_D_ for each benchmark system. As mentioned above, converged state populations and thereby converged K_D_ values (from the state populations) were attainable only after the application of the haMSM analysis procedure. Importantly, K_D_ values calculated from rate constants are within error of those calculated from state populations using the haMSM procedure.

Our results reveal that the effects of applying the haMSM procedure on the efficiencies of the heavy-weight versions of steady-state and equilibrium WE simulations in calculating all the observables of interest varied considerably depending on the number of bins used for the procedure (Fig. 6). The use of the minimal set of 3 bins resulted in similar efficiencies to those from direct calculation of the observables. While in some cases increasing the number of bins increased the efficiency of the haMSM procedure (*e.g*., in calculating k_off_ for the CH_4_/CH_4_ system under equilibrium conditions), in other cases, increasing the number of bins reduced the efficiency (*e.g*. in calculating k_off_ for the K+/CE system under steady-state conditions). Such reductions may be due to a smaller number of observed bin-to-bin transitions between successive WE iterations, resulting in large variations in the transition rate matrix and ultimately, greater uncertainty in the rate constants (Figs. S3 and S4). The same conclusions apply to the light-weight versions of both steady-state and equilibrium WE simulations (Figs. S5 and S6).

**FIG. 6.**
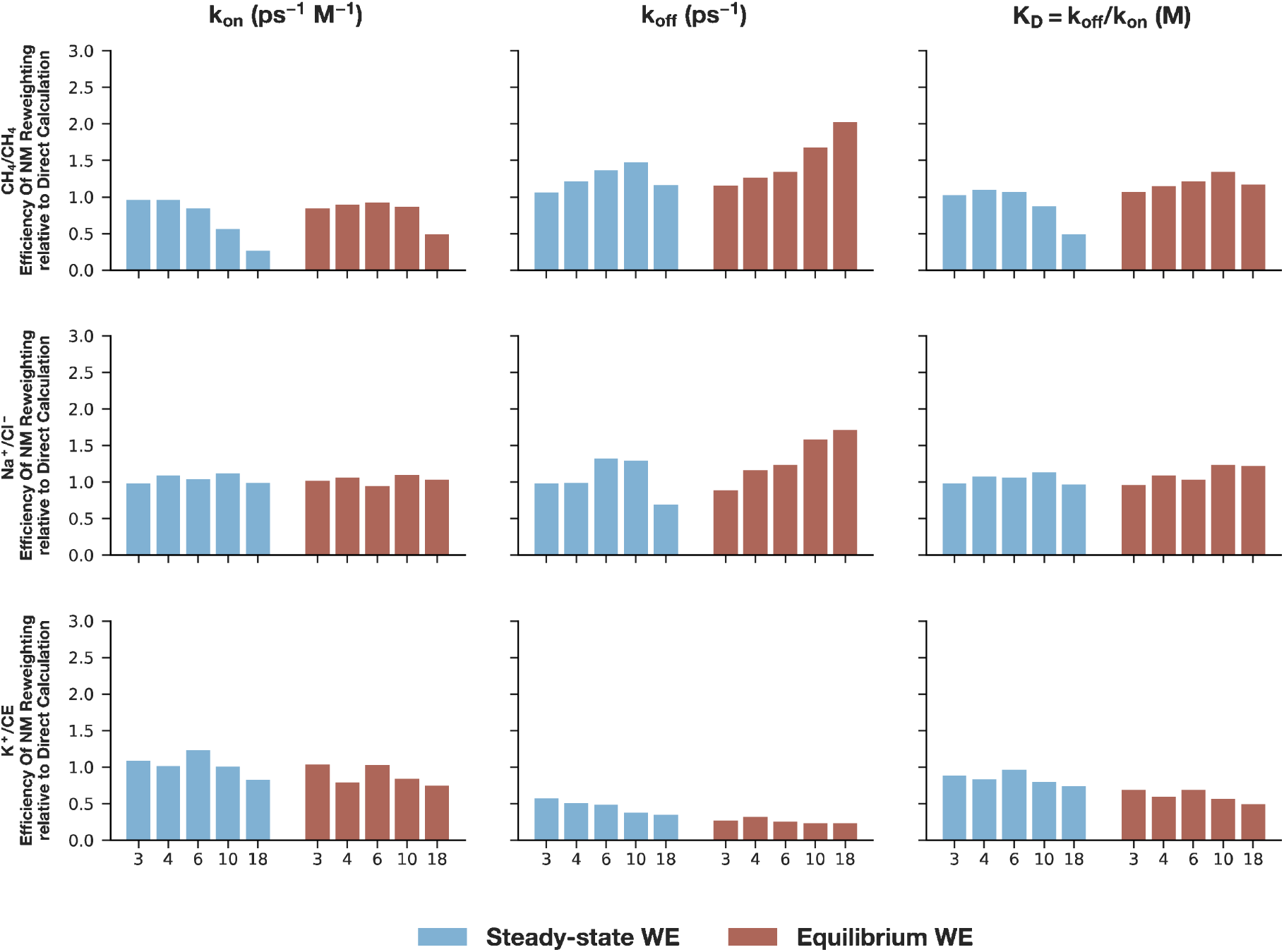
Effect of applying the history augmented Markov State Model [haMSM; previously referred to as non-Markovian (NM) reweighting] post-reweighting procedure on the efficiency of calculating the k_on_, k_off_, and K_D_ for each benchmark system. Data shown is from heavy-weight simulations, both as single equilibrium WE simulations (red) and pairs of steady state WE simulations (blue). Different sets of bins were tested in the application of the haMSM procedure, ranging from a minimal set (3 bins) to 18 bins. The horizontal gray line indicates equal efficiency relative to directly calculating the simulation observables.

For the minimal set of bins, provided that the target states are known in advance, the calculation of K_D_ values from either rate constants or state populations was more efficient using pairs of steady-state WE simulations rather than single equilibrium WE simulations (Fig. 4). Interestingly, the efficiencies of using either steady-state or equilibrium WE simulations in calculating K_D_ values from rate constants vs. state populations are similar for all of the benchmark systems except for the CH_4_/CH_4_ system. For example, pairs of steady-state WE simulations of the CH_4_/CH_4_ system were 3-fold more efficient than brute force simulations in calculating the K_D_ from the directly calculated rate constants, whereas after application of the haMSM post-simulation analysis to both the brute force and steady-state WE simulations calculating the K_D_ from the state populations is 124-fold more efficient (Figs. 2 and 3). This 46-fold difference in efficiency may be due to an insufficient number of observed transitions in the brute force simulations thereby resulting in a large uncertainty in the K_D_ based on state populations (Figs. S1 and S2).

### 4.5 Tracking of trajectory history during equilibrium WE simulations

Finally, we tested whether including the “history” of each trajectory (*i.e.* state last visited) as a separate dimension of the progress coordinate during dynamics propagation would enhance the efficiency of calculating the k_on_, k_off_, and K_D_ using equilibrium WE simulations. As shown in Fig. 7, history tracking during dynamics propagation substantially increased the efficiency of calculating the observables of interest for all of the benchmark systems, with the exception of k_on_ for the Na^+^/Cl^-^ system. The largest efficiency increases were observed for the K^+^/CE system, with 22-, 7-, and 89-fold gains in the efficiencies of calculating the k_off_, k_on_, and K_D_, respectively, relative to brute force simulations. For the CH_4_/CH_4_ and Na^+^/Cl^-^ systems, the efficiencies in calculating the K_D_, both from rate constants and state populations, were either equal to or greater than that of brute force simulations. The target number of trajectories/bin was the same for each history-tracked bin, and as such the simulation spent roughly equal time in both association and dissociation ensembles. Therefore, the efficiency reductions in calculating the k_on_ for both the CH_4_/CH_4_ and Na^+^/Cl^-^ systems may have allowed for the k_off_ to be calculated more efficiently.

**FIG. 7.**
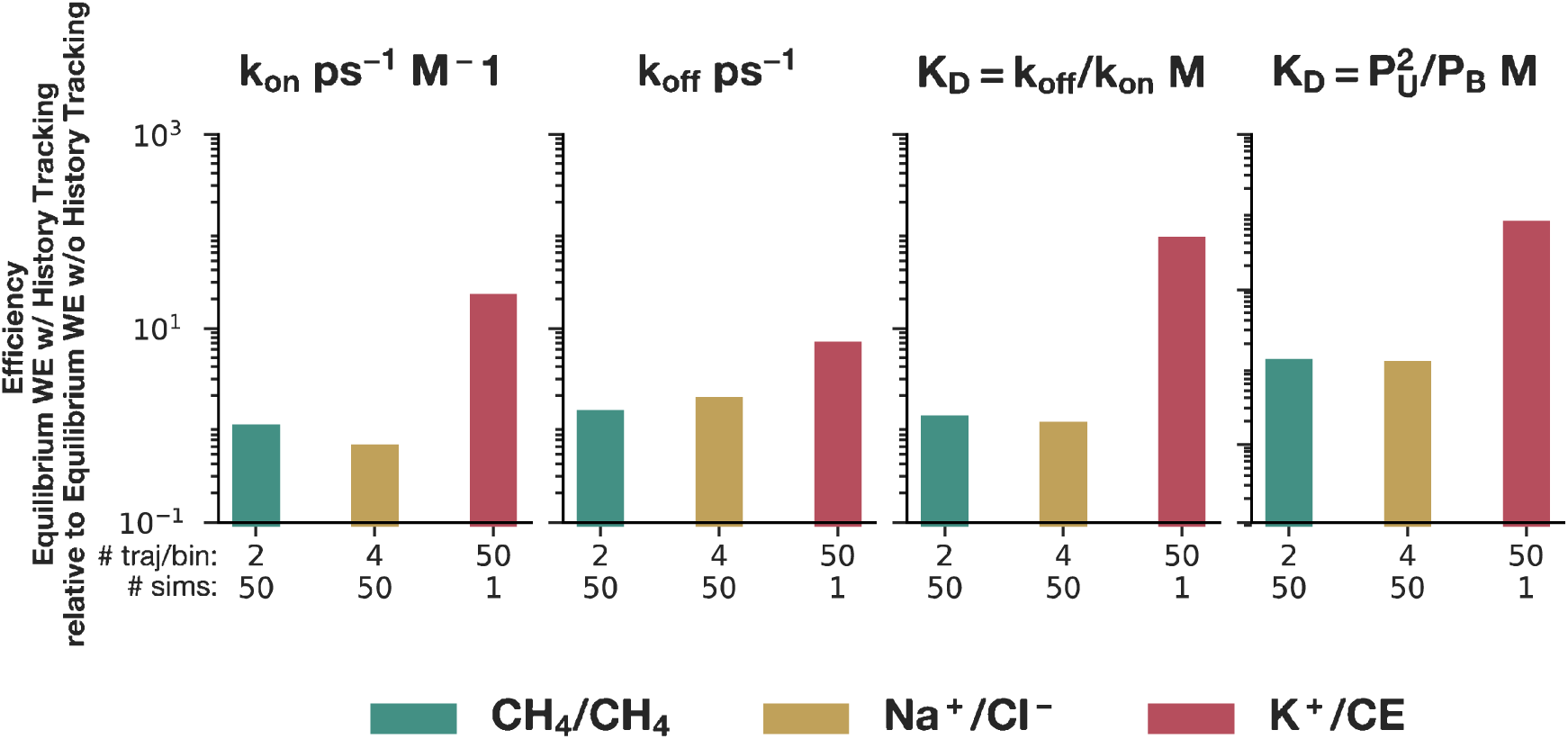
Effect of including trajectory history as part of the progress coordinate during dynamics propagation on the efficiency of equilibrium WE simulations in calculating the k_on_, k_off_, K_D_ for each benchmark system. Only heavy-weight simulations were used in this comparison. The horizontal gray line indicates equal efficiency relative to the same simulations without the inclusion of trajectory history as part of the progress coordinate.

### 4.6 Which WE protocol should you use for your simulation of interest?

As illustrated in Fig. 8, the appropriate WE protocol depends on the goals of the simulations involving the rare event of interest. Furthermore, although some of the advances to the WE strategy were found to reduce the efficiency of calculating certain observables relative to the original WE strategy,^1^ each advance provides appealing features. Our specific recommendations for the most common simulation goals are presented below.

**FIG. 8.**
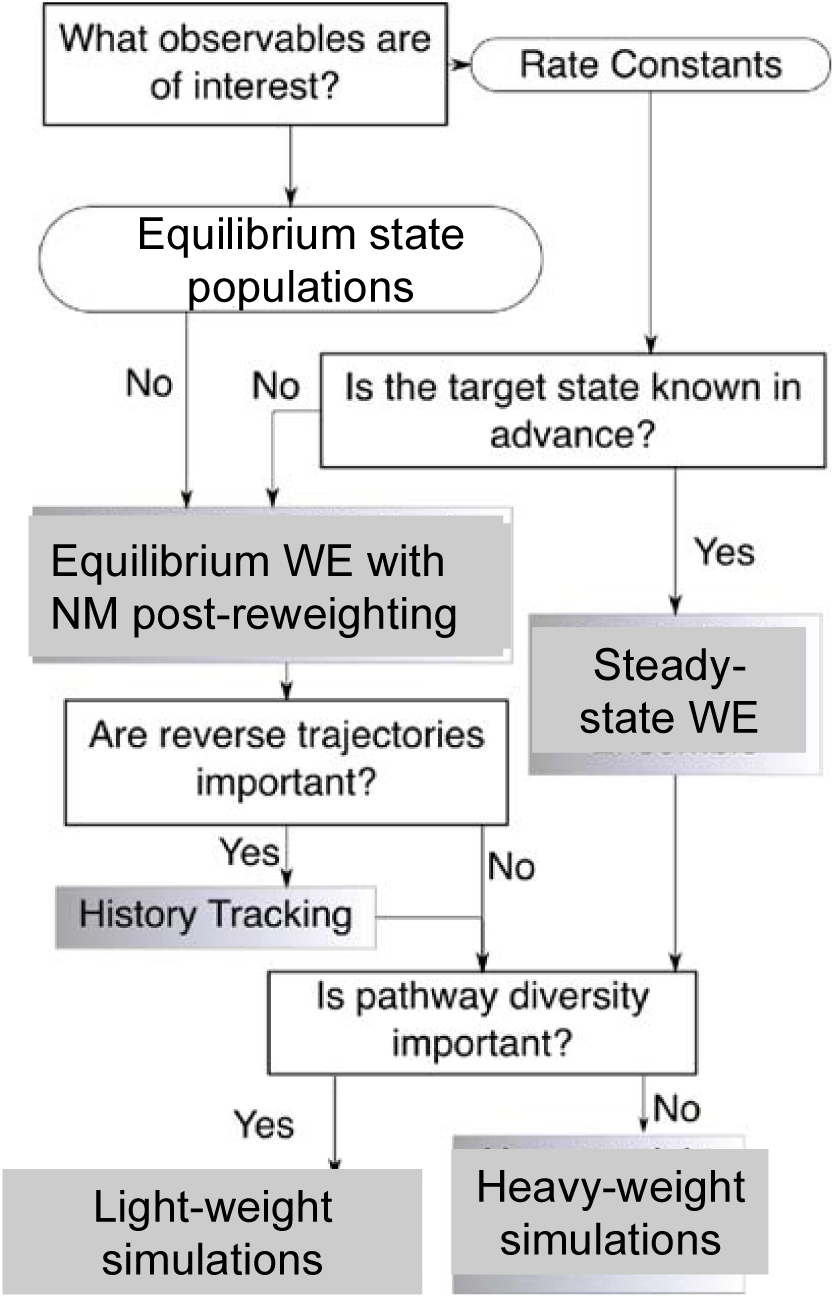
Flowchart for determining a suitable WE simulation protocol for an application of interest.

If the target state of the rare event is well-defined *a priori*, steady state WE simulations are recommended over equilibrium WE simulations for calculating both non-equilibrium and equilibrium observables (*e.g*. rate constants and equilibrium state populations, respectively). In particular, we have demonstrated that rate constants for both the forward and reverse directions of a rare event, as well as the K_D_ values resulting from the ratio of the rare constants, are more efficiently calculated using a pair of steady-state WE simulations in opposite directions rather than a single equilibrium WE simulation. Furthermore, we have demonstrated for the first time that the application of the haMSM post-simulation analysis can enhance the efficiency of the K_D_ from equilibrium state populations. While the target state must be defined in advance for steady state WE simulations, we note that such simulations can be run using a particularly strict (yet still attainable definition) of the target state for the recycling of trajectories, and that this state definition can then be refined to less strict definitions after the completion of the simulations.

If the target state of the rare event is not strictly defined in advance, only equilibrium WE simulations can be carried out since it is not possible to recycle trajectories at the target state to maintain steady state conditions. Although equilibrium WE simulations are not as efficient as steady state WE simulations in calculating rate constants and binding affinities, equilibrium WE simulations are still more efficient than brute force simulations for systems with sufficiently high free energy barriers (*e.g.* K+/CE) and provide the maximum flexibility in refining state definitions after the completion of simulations. As mentioned above, the efficiency of equilibrium WE simulations in calculating the observables of interest can be further enhanced by including the trajectory history (state last visited) as a dimension of the progress coordinate during dynamics propagation in equilibrium WE simulations. This tracking of trajectory history not only ensures the survival of pathways in the opposite direction from the target state, but can increase the diversity of pathways in both directions.

If the primary goal is to calculate rate constants and equilibrium observables (*e.g*. state populations) along with the generation of continuous pathways for the rare event of interest, the use of single heavy-weight simulations is recommended over a large set of light-weight simulations, regardless of whether the simulations are run under steady state or equilibrium conditions. Our results demonstrate that steady-state, heavy-weight simulations are the most efficient strategy for calculating rate constants, particularly for challenging, complex processes such as protein-ligand unbinding where the timescale of the process is far beyond what is accessible to brute force simulations. To calculate equilibrium observables, we recommend applying the haMSM post-simulation analysis to either single equilibrium WE simulations or pairs of steady-state WE simulations in opposite directions.

Notably, we have demonstrated for the first time that the application of the haMSM analysis procedure can enhance the efficiency of computing binding affinities from state populations, provided that the stable states are carefully defined such that the surface of state A from which A-type trajectories are initiated is the same surface at which B-type trajectories end, and vice versa.^18^ We note that the minimal set of bins with the haMSM procedure results in similar efficiencies in the calculation of rate constants as direct calculations and that the use of a greater number of bins may increase this efficiency. However, binning too finely may reduce the efficiency of the calculation. We therefore recommend that different numbers of bins be tested in the application of the haMSM procedure – ranging from the minimal set to the maximum, affordable number of bins – to determine the set of bins that yields the best efficiency in calculating the observable of interest (*e.g.*, generating plots similar to those in Fig. 6).

If the goal is to generate a diverse ensemble of pathways for the rare event of interest, a large set of light-weight simulations may be preferable to a single heavy-weight simulation, provided that the light-weight simulations are carried out using a minimum of 4 target trajectories/bin. This minimum number of target trajectories/bin is required to calculate the observables of interest with greater efficiency than brute force simulations and may vary with the complexity and timescale of the rare event. In principle, single heavy-weight simulations could be combined with improved schemes for the replication and pruning of trajectories to increase the diversity of pathways while leveraging the greater statistical ratcheting that results from the large target number of trajectories/bin. Thus, it is likely that WE protocols will be available in the near future for generating the desired diversity of pathways without sacrificing efficiency in computing both rate constants and equilibrium observables of interest.

Finally, consistent with a previous WE study,^18^ we have found that the calculation of rate constants from a WE simulation can be sensitive to state definitions. For example, equilibrium WE simulations of the K^+^/CE system yield computed k_on_ values that are inconsistent with those from brute force simulations when the unbound state was defined to have a distance of 11.6 Å or 25 Å between the K^+^ ion and center-of-mass of the crown ether oxygens; these distances were chosen from the probability distribution shown in Fig. 1 and correspond to the entrance (inflection point) and bottom of the unfolded state basin, respectively. On the other hand, if a distance of 24 Å is used to define the unbound state, the computed k_on_ values are not only within error of those from brute force simulations, but achieve steady values more quickly than rate constants computed using the other two unbound state definitions. The development of schemes to identify robust state definitions is beyond the scope of this work and would be a valuable future direction for WE strategies. In the meantime, for any given WE simulation, we recommend testing the sensitivity of rate calculations to the state definitions.

## 5. CONCLUSION

We evaluated the impacts of several advances in WE path sampling strategies on the efficiencies of calculating the k_on_, k_off_, and K_D_ for three benchmark systems, listed in order of timescales for association/dissociation: CH_4_/CH_4_, Na^+^/Cl^-^, and K^+^/CE. In particular, we quantitatively assessed the following advances: (i) carrying out a large set of light-weight simulations vs. a single heavy-weight simulation, (ii) the use of equilibrium vs. steady-state WE simulations, (iii) tracking the trajectory history during the dynamics propagation of equilibrium WE simulations, and (iv) history augmented Markov State Model (haMSM) post-simulation analysis of an equilibrium set of trajectories.^7^ It is worth noting that it is just as valuable to determine what does not work as what does work in enhancing the efficiency of calculating the observables of interest. We report the following novel findings.

First, we have demonstrated that the application of the haMSM post-simulation analysis to pairs of steady-state WE simulations in opposite directions can efficiently yield equilibrium observables, *i.e*. K_D_ values. On the other hand, if the target state is not well-defined in advance thereby making it impossible to run steady-state WE simulations, heavy-weight, equilibrium WE simulations with the application of the haMSM analysis procedure are still more efficient than brute force simulations. The application of the haMSM procedure was particularly effective in increasing the efficiency of calculating equilibrium state populations thereby increasing the efficiency of calculating the K_D_ from the state populations. Depending on the number of bins used in the haMSM post-simulation analysis, the analysis could also increase the efficiency of calculating the rate constants, particularly the k_off_ for the most challenging system, K^+^/CE.

Second, our results reveal that a set of 50 light-weight simulations is less effective than a single heavy-weight simulation with 50 target trajectories/bin in calculating the k_on_, k_off_, and K_D_ due to a smaller extent of statistical ratcheting, which is a hallmark of WE strategies. However, light-weight simulations may be preferable for increasing the diversity of pathways by generating a larger number of pathways that are not correlated in history, *i.e.* sharing no common trajectory segments. Alternatively, it may be possible to combine the use of single heavy-weight simulations with improved schemes for replication and pruning of trajectories to both increase the diversity of pathways while maintaining the efficiency associated with heavy-weight simulations in calculating the observables of interest.

Third, we have tested for the first time the incorporation of trajectory history as a dimension of the progress coordinate during dynamics propagation to ensure the generation of trajectories in both the dissociation and association directions. Our results reveal that history tracking can in general, increase the efficiency of calculating the k_on_, k_off_, and K_D_ for the benchmark systems. We note that the incorporation of trajectory history requires definitions of the initial and target states before carrying out the simulation. In cases where states are well-defined in advance, a more efficient simulation protocol than the use of equilibrium WE simulations with history tracking is to run two sets of steady-state WE simulations in opposite directions such that the simulations can be combined to yield an equilibrium set of trajectories to compute both equilibrium and non-equilibrium observables. As the probability of pruning reverse trajectories is determined by the scheme for replicating and pruning trajectories, modifications of this scheme may improve the efficiency of calculating the observables of interest while ensuring the survival of trajectories in the direction that is opposite from the target states.

Among all of the WE protocols tested, steady-state, heavy-weight simulations were found to be the most efficient in calculating the k_on_, k_off_, and K_D_ for the most challenging benchmark system, K^+^/CE. Consistent with this finding is the fact that our previous WE simulations of protein-peptide association involved a heavy-weight protocol under essentially steady-state conditions since pathways were generated primarily for the association process rather than both association and dissociation processes.^6^ For long-timescale processes (microseconds or beyond) for which the target state is well-defined in advance of the simulation, we therefore recommend carrying out heavy-weight, steady-state WE simulations to compute rate constants as well as equilibrium observables (via the combination of steady-state simulations in opposite directions).

Given the unprecedented amount of simulation that was completed for each of the benchmark systems (83.17 μs of aggregate simulation time), the resulting set of simulations provide a valuable reference data set for evaluating future advances to enhanced sampling strategies that maintain rigorous kinetics, including WE strategies with advances that may enhance the diversity of pathways without reducing the efficiency of computing observables of interest.

## SUPPLEMENTARY MATERIAL

WE parameters and conformational sampling simulation details for benchmark systems.

## ACKNOWLEDGMENTS

The authors thank Alex DeGrave, Ali Sinan Saglam, and Karl Debiec for their insightful discussions. This work was supported by NSF CAREER award MCB-0846216 to LTC; NIH grant 1RO1GM115807 to LTC and DMZ. Computational resources were provided by NSF MRI award CNS-1229064 and by the University of Pittsburgh Center for Research Computing. This project has been funded in whole or in part with Federal funds from the National Cancer Institute, National Institutes of Health, under Contract No. HHSN261200800001E. The content of this publication does not necessarily reflect the views or policies of the Department of Health and Human Services, nor does mention of trade names, commercial products, or organizations imply endorsement by the U.S. Government. This research was supported (in part) by the National Institutes of Health.

