## Supplemental Figures S1-S6 and Tables S1-S3 for "Extensive Evaluation of Weighted Ensemble Strategies for Calculating Rate Constants and Binding Affinities of Molecular Association/Dissociation Processes"

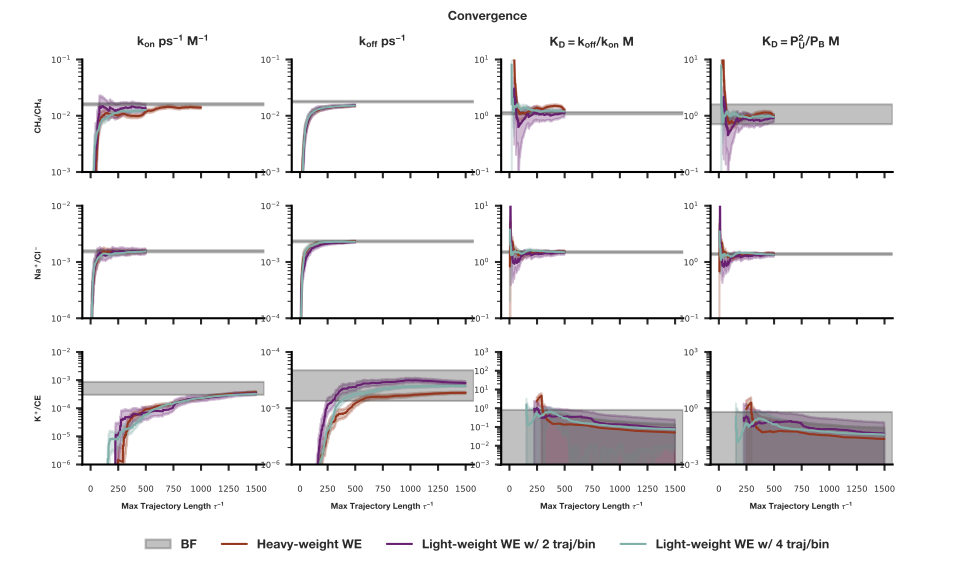
FIG. S1. Evolution of the k_on_, k_off_, and K_D_ to of steady-state WE simulations for each benchmark system. The K_D_ was calculated from both rate constants and state populations; the latter were calculated using the NM analysis procedure. Heavy-weight simulations were carried out with 50 target trajectories/bin and light-weight simulations, with 2 or 4 target trajectories/bin. The gray shaded region represents the average observable and its 95% CI computed from brute force simulations.

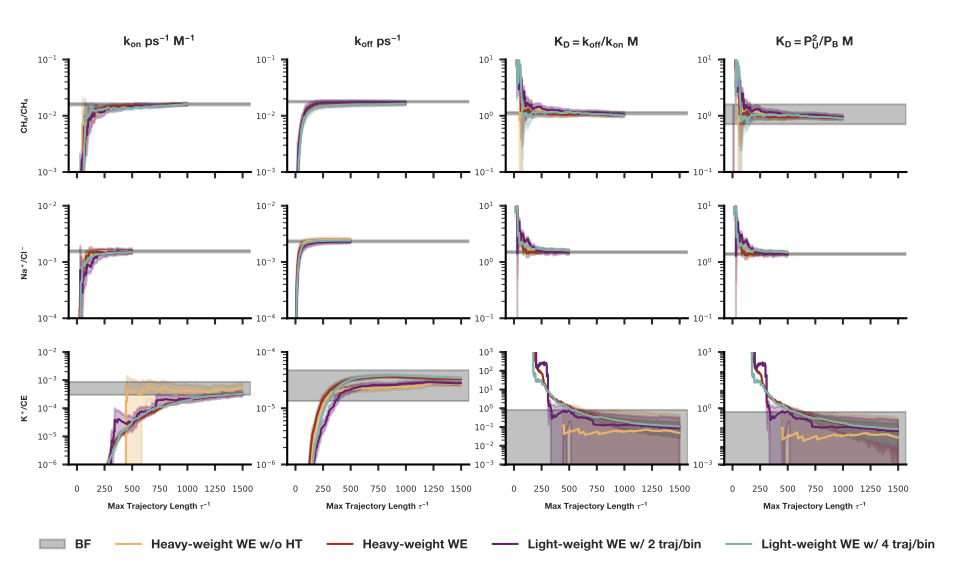
FIG. S2. Evolution of the k_on_, k_off_, and K_D_ of equilibrium WE simulations carried out with history tracking (HT) during dynamics propagation for each benchmark system. The K_D_ was calculated from both rate constants and state populations; the latter were reweighted using an NM procedure. Heavy-weight simulations were carried out with 50 target trajectories/bin and light-weight simulations, with 2 or 4 target trajectories/bin. In addition, a set of heavy-weight simulations were run without history tracking (HT) during dynamics propagation. The gray shaded region represents the average observable and its 95% CI computed from brute force simulations.

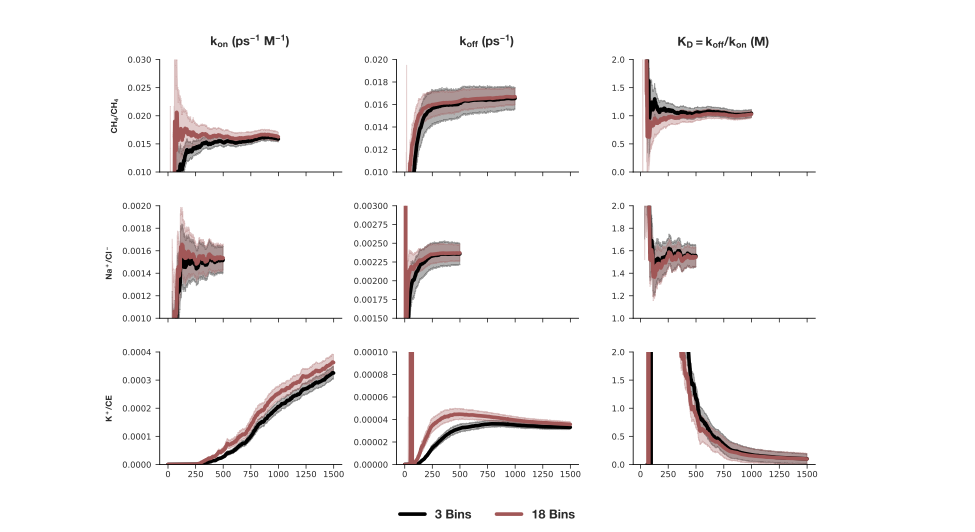
FIG. S3. Effect of varying the number of bins in the application of the NM post-simulation analysis procedure on the convergence of the average k_on_, k_off_, and K_D_ for each benchmark system. Data shown is from equilibrium, heavy-weight simulations. The effect of increasing the number of bins on the efficiencies in calculating the observables of interest can depend strongly on the uncertainty in the observable, *e.g.* the uncertainty (95% CI) of the k_off_ for the CH_4_/CH_4_ system is reduced by the NM procedure, resulting in an increased efficiency in calculating the k_off_, whereas the increased uncertainty of the k_off_ for the K^+^/CE system results in a reduced efficiency in calculating the k_off_.

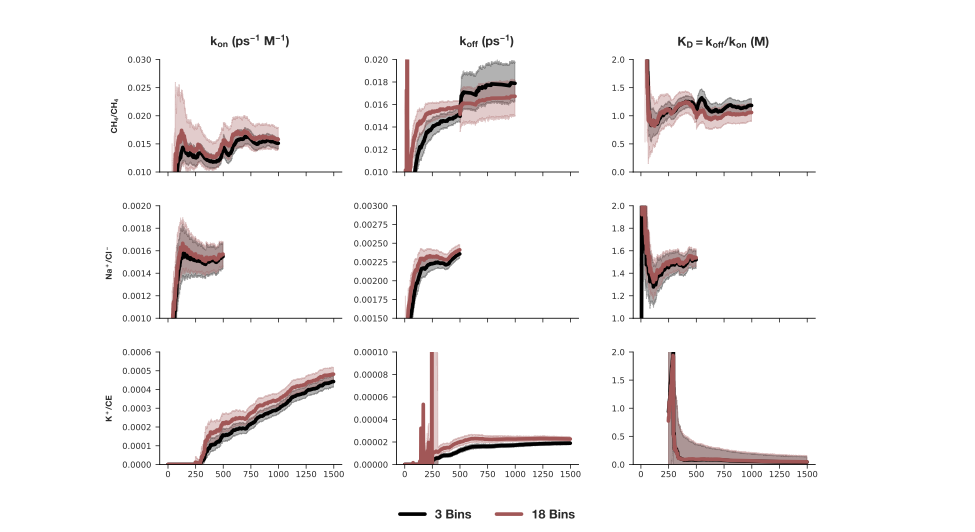
FIG. S4. Effect of varying the number of bins in the application of the NM post-simulation analysis on the convergence of the average k_on_, k_off_, and K_D_ for each benchmark system. Data shown is from steady-state, heavy-weight simulations. The effect of increasing the number of bins on the efficiencies in calculating the observables of interest can depend strongly on the uncertainty in the observable, *e.g.* the uncertainty (95% CI) of the k_off_ for the Na^+^/Cl^-^ system is reduced by the NM analysis procedure, resulting in an increased efficiency in calculating the k_off_, whereas the uncertainty of the k_on_ for the Na^+^/Cl^-^ system is unaffected, resulting in no change in the efficiency in calculating the k_on_.

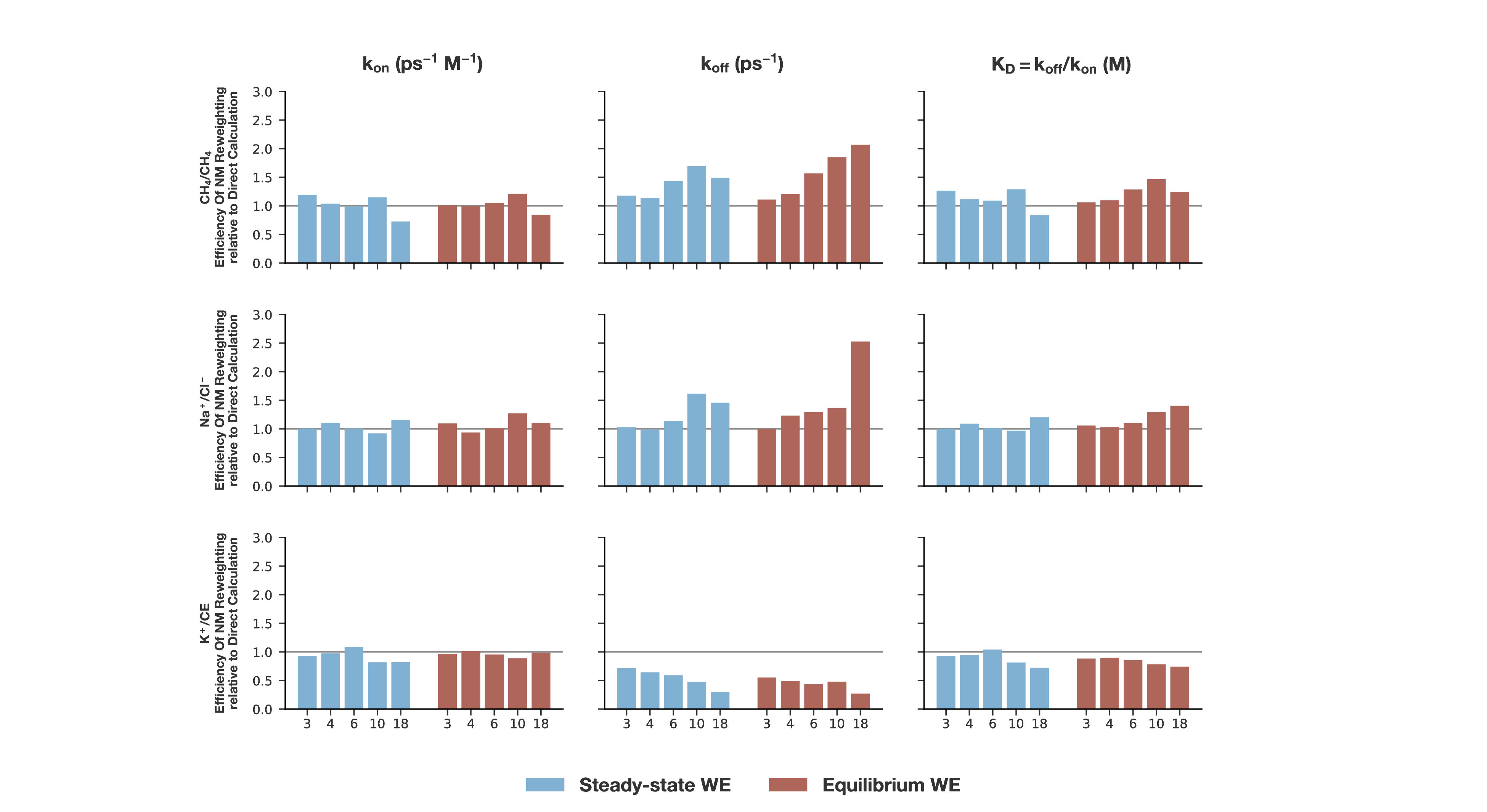
FIG. S5. Effect of applying the non-Markovian (NM) post-simulation analysis on the efficiency of calculating the k_on_, k_off_, and K_D_ for each benchmark system. Data shown is from light-weight simulations with 2 target trajectories/bin, both as a set of 50 equilibrium WE simulations (red) and a set of 100 steady state WE simulations (blue). Different sets of bins were tested in the application of the NM procedure, ranging from a minimal set (3 bins) to 18 bins. The horizontal gray line indicates equal efficiency relative to directly calculating the simulation observables.

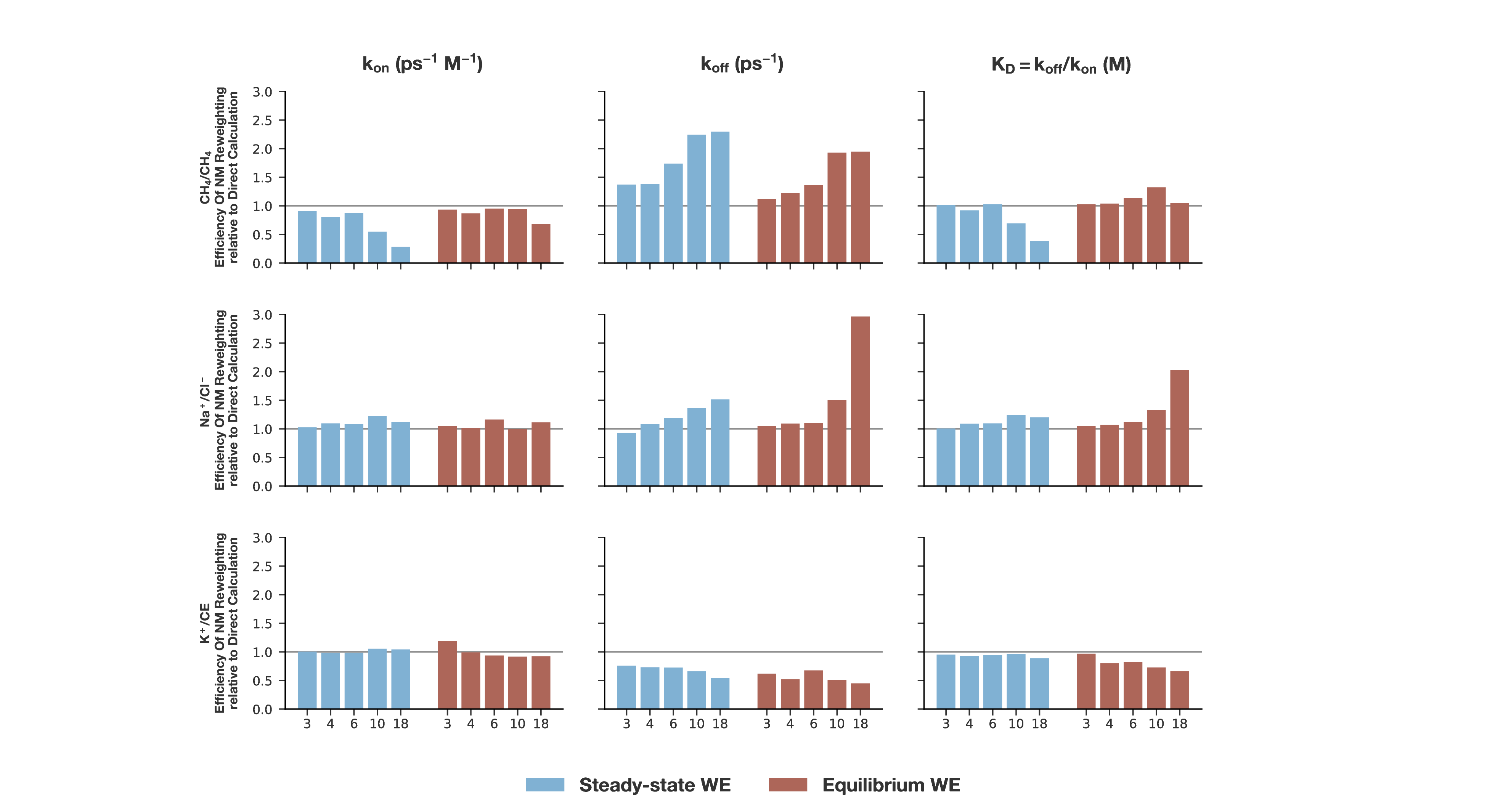
FIG. S6. Effect of applying the non-Markovian (NM) post-simulation analysis on the efficiency of calculating the k_on_, k_off_, and K_D_ for each benchmark system. Data shown is from light-weight simulations with 4 target trajectories/bin, both as a set of 50 equilibrium WE simulations (red) and a set of 100 steady state WE simulations (blue). Different sets of bins were tested in the application of the NM procedure, ranging from a minimal set (3 bins) to 18 bins. The horizontal gray line indicates equal efficiency relative to directly calculating the simulation observables.

**Table S1**. Number of initial-state conformations generated for WE simulations of molecular association/dissociation processes of each benchmark system. Conformations were pre-equilibrated and generated using equilibrium WE simulations.

| system | # bound-state conformations | # unbound-state conformations |
| --- | --- | --- |
| CH_4_/CH_4_ | 19 | 420 |
| Na^+^/Cl^-^ | 98 | 186 |
| K^+^/CE | 293 | 112 |

**Table S2.** State definitions for simulations of benchmark systems. State definitions are similar to those used in Zwier, et al.,^1^ but have been modified for CH_4_/CH_4_ and K+/CE to capture the entirety of the bound state well and the narrowest possible region as the unbound state, similar to the state definitions for Na+/Cl-.

| System | bound state (Å) | unbound state (Å) |
| --- | --- | --- |
| CH_4_/CH_4_ | 4.99 | 14.98 |
| Na^+^/Cl^-^ | 2.80 | 14.98 |
| K^+^/CE | 2.19 | 24.00 |

**Table S3:** Bin boundaries (r/Å) for WE simulations of benchmark systems. Bin boundaries for association are similar to those from Zwier *et al*. (ref), with modifications made for the dissociation process to ensure smooth unbinding for steady state simulations. Bin boundaries for equilibrium simulations are the joint set of both steady state directions.

| CH_4_/CH_4_ | | | Na^+^/Cl^-^ | | | K^+^/18-Crown-6-Ether | | |
| --- | --- | --- | --- | --- | --- | --- | --- | --- |
| EQ | Dissoc. | Assoc. | EQ | Dissoc. | Assoc. | EQ | Dissoc. | Assoc. |
| 4.0  4.75  5.50  6.25  7.00  7.75  8.50  9.25  10.00  11.00  12.00  13.00  14.00  14.98 | 4.0  4.75  5.50  6.25  7.00  7.75  8.50  9.25  10.00  11.00  12.00  13.00  14.00  14.98 | 4.0  4.75  5.50  6.25  7.00  7.75  8.50  9.25  10.00 | 2.80  2.88  3.00  3.10  3.29  3.79  3.94  4.12  4.39  5.43  5.90  6.90  7.90  8.90  9.90  10.90  11.90  12.90  13.90  14.90  15.90 | 2.80  2.88  3.00  3.10  3.29  3.79  3.94  4.12  4.39  5.43  5.90  6.90  7.90  8.90  9.90  10.90  11.90  12.90  13.90  14.90 | 2.80  2.88  3.00  3.10  3.29  3.79  3.94  4.12  4.39  5.43  5.90  6.90  7.90  8.90  9.90  10.90  11.90  12.90  13.90  14.90 | 0.2  0.7  1.5  2.5  3.8  5.0  6.0  7.0  8.0  9.0  10.0  11.6  12.0  13  14  15  16  17  18  19  20  21  22 | 0.2  0.7  1.5  2.5  3.8  5.0  6.0  7.0  8.0  9.0  10.0  11.6  12.0  13.0  14.0  15.0  16.0  17.0 | 1.30  2.5  3.8  5.0  6.0  7.0  8.0  9.0  10.0  11.6  12.0  13  14  15  16  17  18  19  20  21  22 |
